## Supplemental tables and survey for "Biomedical researchers’ perspectives on the reproducibility of research: a cross-sectional international survey"

**Supplementary Materials.**

**S1. Search strategy**

We will obtain a list of all journals indexed in MEDLINE together with their NLM ID. A random list of 1,000 journals will be generated in Excel using the RAND() function.

**Two potential strategies can be used to retrieve the articles:**

1. We will search for all articles published in each journal using the search strategy *1234567.jc.* where “1234567” is the NLM ID of the journal. Where NLM ID is not available, we will use the syntax “Name of journal”.nj.

We will run search for each journal separately. After each search, we will sort the results by Entry date (descending) and export the first 20 results.

Pros: All journals are equally represented.

Cons: Time consuming, as the results need to be exported manually for each journal

2. We will carry out a combined search for all journals using the following search strategy:
1. 1234567.jc.
2. 2345678.jc.
3. 3456789.jc.
4. or/1-3
5. limit 4 to dt=yyyymmdd-yyyymmdd

Pros: Faster, potentially represent the distribution of articles among journals

Cons: Unequal number of articles per journal; may end up with >20,000 articles (some journals have more than others within the same time frame), we can tweak the date range to get as close to 20,000 as possible.

**Three potential strategies can be used to obtain the email addresses of authors:**

1. All retrieved articles will be re-imported to EndNote/Zotero/Mendeley to retrieve PMID numbers. The list of PMID numbers will be exported as an .csv file and input into an R script (built based on the easyPubMed package) to retrieve the authors’ name, affiliation institutions and email addresses.

Pros: Quick & reliable results, retrieve multiple email addresses

Cons: Not working if the PubMed page does not display email addresses (usually for newly-indexed studies – we can tweak the date rate to mitigate this)

2. In addition, we will use the Find Full Text function in EndNote to retrieve PDF files of these articles, and run these files in another R script for text recognition to extract email addresses.

Pros: High rates of success

Cons: Only retrieve one email address, only work if PDF is present, sometimes false positives (e.g. publisher email address)

3. Any articles where email addresses cannot be retrieved from both methods will be manually screened.

Results from all three methods will be combined into the final list and counter-checked by another author for potential errors before survey distribution.

**S2. Study survey**

**Demographics**

1. What describes you best?

Graduate student

Postdoctoral fellow

Faculty member/PI

Research support staff (E.g., research manager, research associate, technician)

Scientist in industry

Scientist in third sector (E.g., NGO, non-profit)

Government scientist

Other, please specify

2. What is your gender?
   a. Female
   b. Male
   c. Non-binary
   d. Prefer to self-describe:
   e. Prefer not to say
3. What country are you currently employed in? (drop down list)
4. Which of the following best describes your research area?
   a. Clinical research
   b. Preclinical research – in vivo
   c. Preclinical research – in vitro
   d. Health systems research
   e. Methods research
   f. Other, please specify

**Reproducibility perceptions**

For the purposes of this survey, we consider a study to be reproduced when its findings are confirmed in similar experimental systems (these may include slight variations in methods or materials.)  By contrast, a study is replicated when it is repeated exactly. This survey talks about the larger issue of reproducibility of results, not just replication.

5. In your view, is there a reproducibility crisis in biomedicine?
   a. Yes, significant crisis
   b. Yes, a slight crisis
   c. No, there is no crisis
   d. Don’t know
6. What proportion of papers in biomedicine do you think are reproducible?

0% - 100%, in 10% increments.

Biomedicine overall

Clinical biomedical research

In vivo biomedical research

In vitro biomedical research

7. In your view, which of the factors below contribute to irreproducible biomedical research results?

Always contributes; Usually contributes; Sometimes contributes; does not contribute; unsure.

a. Selective reporting of the published literature
b. Pressure to publish
c. Low statistical power
d. Poor statistical analysis
e. Not enough internal replication (E.g., by the original lab/authors)
f. Insufficient study oversight
g. Lack of training in reproducibility
h. Failure to make materials openly available
i. Failure to make original study data openly available
j. Poor study design
k. Fraud
l. Poor quality peer review
m. Problems in the design of replication studies
n. Technical expertise required for replication
o. Variability of standard reagents
p. Bad luck
q. Other, please specify

**Reproducibility experiences**

8. Have you ever tried to replicate a published study YOU previously conducted and failed (i.e., re-ran an experiment but got different results from the original study)?
   a. Yes
   b. No- all replications I have completed of my own research have been successful
   c. No – I have never tried to replicate my own research

If a or b;

A. Did you publish your replication study results? Note: if you have conducted more than one replication study of your own work please respond based on your most recent study.
   a. Yes – but it took longer to publish than other papers you’ve published that were not replications
   b. Yes – and it took about the same amount of time to publish as other papers you’ve published that were not replications
   c. Yes – but it was quicker to publish than other papers you’ve published that were not replications
   d. No – I have submitted but not yet had the work accepted
   e. No – I have not yet submitted, but intend to do so
   f. No – I don’t intend to attempt to publish this study
   g. No- Journals don’t appear interested in publishing replications
   h. Other, please specify
B. What was your motivation for replicating your own study?
9. Have you ever tried to replicate a published study conducted by another team of authors and failed?
   a. Yes
   b. No- all replications I have completed have been successful
   c. No – some [< 100%] of the replications have been successful
   d. No – I have never tried to replicate someone else’s published research

If a or b;

A. Did you publish your replication study results? Note: if you have conducted more than one replication study of another groups research, please respond based on your most recent study.

a. Yes – but it took longer to publish than other papers you’ve published that were not replications
b. Yes – and it took about the same amount of time to publish as other papers you’ve published that were not replications
c. Yes – but it was quicker to publish than other papers you’ve published that were not replications
d. No – I have submitted but not yet had the work accepted
e. No – I have not yet submitted, but intend to do so
f. No – I don’t intend to attempt to publish this study
g. Other, please specify

B. What was your motivation for replicating another researcher’s study?
10. Have you ever been contacted by another researcher who was unable to reproduce a finding you published?
    a. Yes
    b. No
    c. Unsure

**Reproducibility support**

11. Does your research institution have established procedures to enhance reproducibility of biomedical research?
    a. Yes
    b. No
    c. Unsure

If yes, please describe what processes are in place to enhance reproducibility:

12. My institution would value me doing new biomedical studies more than me doing replication studies.
    a. True
    b. False
    c. Unsure
13. In my biomedical research setting it would be harder to find funding to conduct a replication study than it would be to find funding for a new study.
    a. True
    b. False
    c. Unsure
14. Are you aware of funders providing specific calls for conducting reproducibility related research (e.g., to reproduce studies, to conduct meta-science on reproducibility)?
a. Yes ; If so, which ones
b. No
15. Does your research institution provide training on how to enhance the reproducibility of research?
a. Yes, and I have taken it ; please specify
b. Yes, but I have not taken it
c. No.
d. Unsure
16. Do you think your views on reproducibility represent those of the average researcher in your field at your career stage?
a. Yes
b. No
17. Is there anything else about reproducibility you would like to share with us?

(open text)

S3. Additional analyses of survey responses to reproducibility perception items by gender. Note that we present data for males and females only; response values for other gender options were low (all n<3)

| **Item** | **Response options** | **Male**  **(N=943)** | | **Female (N=643)** | |
| --- | --- | --- | --- | --- | --- |
|  |  | **N** | **%** |  |  |
| In your view, is there a reproducibility crisis in biomedicine? | Yes, a significant crisis | 259 | 28 | 151 | 24 |
|  | Yes, a slight crisis | 437 | 46 | 283 | 44 |
|  | No, there is no crisis | 155 | 16 | 82 | 13 |
|  | Don’t know | 92 | 10 | 127 | 20 |
|  | *Missing data* | - | - | - | - |

Additional analyses of survey responses to reproducibility perception items by academic role

| **Role** | **In your view, is there a reproducibility crisis in biomedicine?** | **N** | **%** |
| --- | --- | --- | --- |
| Other | Yes, a significant crisis | 19 | 26 |
|  | Yes, a slight crisis | 29 | 40 |
|  | No, there is no crisis | 10 | 14 |
|  | Don’t know | 15 | 21 |
|  | Total | 73 | 100 |
| Graduate student | Yes, a significant crisis | 28 | 32 |
|  | Yes, a slight crisis | 37 | 42 |
|  | No, there is no crisis | 8 | 9 |
|  | Don’t know | 15 | 17 |
|  | Total | 88 | 100 |
| Postdoctoral fellow | Yes, a significant crisis | 31 | 24 |
|  | Yes, a slight crisis | 63 | 49 |
|  | No, there is no crisis | 13 | 10 |
|  | Don’t know | 22 | 17 |
|  | Total | 129 | 100 |
| Faculty member/PI | Yes, a significant crisis | 288 | 25 |
|  | Yes, a slight crisis | 536 | 47 |
|  | No, there is no crisis | 181 | 16 |
|  | Don’t know | 146 | 13 |
|  | Total | 1151 | 100 |
| Research support staff (E.g., research manager, research associate, technician) | Yes, a significant crisis | 16 | 30 |
|  | Yes, a slight crisis | 21 | 39 |
|  | No, there is no crisis | 5 | 9 |
|  | Don’t know | 12 | 22 |
|  | Total | 54 | 100 |
| Scientist in industry | Yes, a significant crisis | 10 | 36 |
|  | Yes, a slight crisis | 12 | 43 |
|  | No, there is no crisis | 5 | 18 |
|  | Don’t know | 1 | 4 |
|  | Total | 28 | 100 |
| Scientist in third sector (E.g., NGO, non-profit) | Yes, a significant crisis | 11 | 41 |
|  | Yes, a slight crisis | 8 | 30 |
|  | No, there is no crisis | 6 | 22 |
|  | Don’t know | 2 | 7 |
|  | Total | 27 | 100 |
| Government scientist | Yes, a significant crisis | 15 | 28 |
|  | Yes, a slight crisis | 22 | 41 |
|  | No, there is no crisis | 9 | 17 |
|  | Don’t know | 8 | 15 |
|  | Total | 54 | 100 |

S4. List of training links provided by participants

| Name/Organization providing training | Link | Format of training | Open status |
| --- | --- | --- | --- |
| CITI program: Biomedical Responsible Conduct of Research, Rigor, Reproducibility and Ethical Behavior in Biomedical Research | <https://about.citiprogram.org/news/improve-study-design-to-promote-reproducibility/> | Online course | No |
| Universidade de Sao Paulo - Ribeirao Preto College of Nursing | <http://www.eerp.usp.br/research-home/> | University resources | Unclear |
| Universite de Montreal - The reproducibility of research: Issues and good practices | <https://calendrier.bib.umontreal.ca/event/3617795> | Standalone lecture | Unclear |
| Duke University School of Medicine - Reproducibility Crisis: What Can We Do? | <https://medschool.duke.edu/events/reproducibility-crisis-what-can-we-do> | University event | Unclear |
| The University of Utah | <https://education.research.utah.edu/rigor.php> | University course | No |
| UC Davis: Training Program in Molecular and Cellular Biology | <https://mcbtrainingprogram.ucdavis.edu/quantitative-approaches-molecular-and-cellular-biology-rigor-and-reproducibility> | University course | No |
| University of Oxford - Integrity and ethics training | <https://researchsupport.admin.ox.ac.uk/support/training/ethics> | University courses and workshops | No |
| Software Carpentry | <https://software-carpentry.org> | In-person and online workshops | Yes |
| Clinical Epidemiology Unit (CEU) All India Institute of Medical Sciences Ansari Nagar, New Delhi - Designing Medical Research Workshop and Thesis | <https://www.aiims.edu/aiims/events/nero_conf.htm> | In-person workshop | Unclear |
| Columbia University Irving Medical Center - Responsible Conduct of Research and Related Policy Issues | <https://www.gsas.cuimc.columbia.edu/responsible-conduct-research-and-related-policy-issues> | University Course | No |
| IECS Institute for Clinical Effectiveness and Health Policy | <https://www.iecs.org.ar/en/> | Online courses | Unclear |
| King's College London Riot Science Club | <https://www.kcl.ac.uk/events/series/riot-science-club> | Seminar series | Yes |
| Memorial Sloan Kettering Cancer Center - Responsible Conduct of Research (RCR) | <https://www.mskcc.org/sites/default/files/node/4368/documents/rtm-1008_rcr_2018updated_0.pdf> | University course | No |
| Radboud umc university medical center | <https://www.radboudumc.nl/en/research/scientific-integrity> | University courses | No |
| Brazilian Reproducibility Initiative | <https://www.reprodutibilidade.bio.br/home> | Online webinars and events | Yes |
| The University of Alabama at Birmingham Center for Clinical and Translational Science | <https://www.uab.edu/ccts/training-academy/kaizen/kaizen-r2t> | University training platform | Yes |
| University College Cork, Ireland - Epigeum Online Research Integrity Training | <https://www.ucc.ie/en/research/support/integrity/researchintegritytraining/epigeumonlineresearchintegritytraining/> | University course | No |
| Karlsruhe Institute of Technology - Theory of Science and Ethics in Biology | <https://www.botanik.kit.edu/botzell/953.php> | University course | No |
| The University of Edinburgh | <https://www.ed.ac.uk/research-office/research-talent-and-culture/research-improvement> | Standalone website resources and online workshops | Yes |
| British Neuroscience Association | <https://www.bnacredibility.org.uk> | Standalone website resources and webinars | Yes |
| Swiss Personalized Health Network incentive | <https://sphn.ch> | Standalone website resources and training events | Yes |
| Society for Neuroscience | <https://www.sfn.org> | Standalone website resources and virtual events | Yes |
| University of Zurich, Center for Reproducible Science | <https://www.crs.uzh.ch/en.html> | University course materials | Yes |
| Aarhus University | <https://medarbejdere.au.dk/en/administration/researchandtalent/responsible-conduct-of-research> | University course | No |
